# Optogenetic stimulation of kisspeptin neurones within the posterodorsal medial amygdala increases LH pulse frequency in female mice

**DOI:** 10.1101/497164

**Authors:** G Lass, XF Li, RA de Burgh, W He, Y Kang, S-H Yeo, Lydia Sinnett-Smith, WH Colledge, MS Grubb, SL Lightman, KT O’Byrne

## Abstract

Kisspeptin within the arcuate nucleus of the hypothalamus is a critical neuropeptide in the regulation of reproduction. Together with neurokinin B and dynorphin A, arcuate kisspeptin provides the oscillatory activity that drives the pulsatile secretion of GnRH, and therefore LH pulses, and is believed to be a central component of the GnRH pulse generator. It is well established that the amygdala also exerts an influence over gonadotrophic hormone secretion and reproductive physiology. The discovery of kisspeptin and its receptor within the posterodorsal medial amygdala (MePD), and our recent finding showing that intra-MePD administration of kisspeptin or a kisspeptin receptor antagonist results in increased LH secretion and decreased LH pulse frequency, respectively, suggests an important role for amygdala kisspeptin signalling in the regulation of the GnRH pulse generator. To further investigate the function of amygdala kisspeptin, the present study used an optogenetic approach to selectively stimulate MePD kisspeptin neurones and examine the effect on pulsatile LH secretion. MePD kisspeptin neurones in conscious Kiss1-CRE mice were virally infected to express a channelrhodopsin protein and selectively stimulated by light via a chronically implanted fibre optic cannula. Continuous stimulation using 5 Hz resulted in an increased LH pulse frequency, which was not observed at the lower stimulation frequencies of 0.5 and 2 Hz. In wild-type animals, continuous stimulation at 5 Hz did not affect LH pulse frequency. These results demonstrate that selective activation of MePD Kiss1 neurons can modulate hypothalamic GnRH pulse generator frequency.

## Introduction

It is well established that hypothalamic kisspeptin (Kp) is a critical neuropeptide for reproduction. Inactivating mutations of the genes encoding *KISS1* or its receptor, *KISS1R (a.k.a. GPR54)*, result in hypogonadotrophic hypogonadism and a failure to progress through puberty in humans and rodent models (1, 2). Of the major hypothalamic populations of *Kiss1* neurones, in the anterior preoptic area and the arcuate nucleus (ARC), attention has focused on the ARC as a key component of the GnRH pulse generator (3). These neurones, known as KNDy, because they co-express neurokinin B (NKB) and dynorphin A (DYN), innervate the distal processes of the GnRH neurones at the level of the median eminence (4). NKB acting on its receptor (TACR3) is thought to function as an excitatory signal to depolarise KNDy cells postsynaptically in the neural network, resulting in kisspeptin output to the GnRH neurones to initiate each GnRH pulse. The co-released DYN functions as an inhibitory signal within the KNDy neural network, acting presynaptically on kappa opioid receptors (KOR) to inhibit the release of NKB, thus terminating kisspeptin release and terminating the signal for GnRH secretion (5).

Despite the autonomous nature of the GnRH pulse generator, it is modulated by various signalling systems, including metabolic, circadian and stress, to regulate reproductive function. The amygdala, a key limbic brain structure commonly known for its role in higher-order emotional processing, is implicated in reproduction, including psychological stress-induced suppression of pulsatile LH secretion (6) and therefore it is reasonable to suggest that the amygdala is a component of upstream regulation of the hypothalamic GnRH pulse generator. The finding of extra-hypothalamic *Kiss1* expression and its receptor in the medial amygdala, and more specifically its posterodorsal subnucleus (MePD) (7), has opened up new possibilities concerning its role in reproductive function. Indeed, we have shown that the MePD *Kiss1* neurones are a key upstream regulator of pubertal timing (8), sexual motivation and social behaviour (9). Moreover, we have discovered through neuropharmacological studies that Kp signalling in the MePD *per se* robustly regulates hypothalamic GnRH pulse generator frequency (10), however, the underlying neural mechanisms are unknown.

In this study, we have used an optogenetic approach to selectively stimulate the MePD *Kiss1* neurones in fully-conscious mice to characterise the parameters that alter LH pulse frequency.

## Materials and methods

### Animals

Adult female mice weighing between 19-23 g were used. They were bred in house at King’s College London and genotyped with a multiplex PCR protocol to detect heterozygosity for the Kiss-Cre or wild-type allele (11). The primers to detect the wild-type allele were mKiss hetF3 (CCG TCA TCC AGC CTA AGT TTC TCA C) and mKiss hetR3 (ATA GGT GGC GAC ACA GAG GAG AAG C), and the primers to detect the mutant allele were mKiss a526 (GCT TTT ATT GCA CAA GTC TAG AAG CTC) and Asc403 (CAG CCG AAC TGT TCG CCA GGC TCA AGG). Mice were kept singularly housed under controlled conditions (12:12 h hour dark/light cycle, on at 07:00 h, 25 °C) and provided with food and water *ad libitum*. All animal procedures performed were approved by the Animal Welfare and Ethical Review Body (AWERB) Committee at King’s College London, and in accordance with the UK Home Office Regulations.

### Stereotaxic injection of channelrhodopsin viral construct and implantation of fibre optic cannula

Stereotaxic injection of the channelrhodopsin (ChR2) virus, AAV9.EF1.dflox.hChR2(H134R)-mCherry.WPRE.hGH (4.35 × 10^13^ GC/ml; Penn Vector Core, University of Pennsylvania, USA), was performed under aseptic conditions. General anaesthesia was achieved using ketamine (Vetalar, 100 mg/kg, i.p.; Pfizer, Sandwich, UK) and xylazine (Rompun, 10 mg/kg, i.p.; Bayer, Leverkusen, Germany). Animals were secured in a motorised Kopf stereotaxic frame and surgical procedures were performed using a robot stereotaxic system (Neurostar, Tubingen, Germany). A small hole was drilled in the skull at a location above the MePD. The stereotaxic injection coordinates used to target the MePD were obtained from the mouse brain atlas of Paxinos and Franklin (12) (2.1 mm lateral, 1.70 mm posterior to bregma and at a depth of 5.1 mm). Using a 2-μL Hamilton micro-syringe (Esslab, Essex, UK) attached to the robot stereotaxic frame, 1 μl of the AAV-construct was injected unilaterally into the MePD at a rate of 100 nl/min. The needle was left in position for a further 5 min and then removed slowly over 1 min. A fibre optic cannula (200 μm, 0.39NA, 1.25 mm ceramic ferrule; Thorlabs LTD, Ely, UK) was then inserted at the same co-ordinates as the injection site, but to a depth of 4.85 mm, so that the fibre optic cannula was situated immediately above the latter. A glue composite (RS Pro 20 g Super Glue, RS Components, Corby, UK) was then used to fix the cannula in place, and the skin incision closed with suture. After surgery, mice were left for 4 weeks to achieve effective opsin expression. Following a 1 week recovery period from surgery, the mice were handled daily to acclimatise them to the tail-tip blood sampling procedure (13).

### *In vivo* optogenetic stimulation of MePD kisspeptin neurones and blood samplings for LH measurement

Prior to optogenetic stimulation, the very tip of the mouse’s tail was excised using a sterile scalpel for subsequent blood sample collection (14). The chronically implanted fibre optic cannula was then attached via a ceramic mating sleeve to a multimode fibre optic rotary joint patch cables (Thorlabs), allowing freedom of movement of the animal, for delivery of blue light (473 nm wavelength) using a Grass SD9B stimulator controlled DPSS laser (Laserglow Technologies, Toronto, Canada). Laser intensity at the tip of the fibre optic patch cable was 5 mW. After 1 h acclimatisation, blood samples (4 µl) were collected every 5 min for 2.5 h. After 1 h controlled blood sampling, continuous optic stimulation (5-ms pulse width) was initiated at 0.5, 2 or 5 Hz for 90 min. Controls received no optic stimulation. Kiss-Cre mice received the stimulation protocols in random order, with at least two day rest between sampling days. Wild-type (n = 2) received 5 Hz optic stimulation only.

The blood samples were processed by ELISA as reported previously (13). Mouse LH standard and antibody were purchased from Harbour-UCLA (California, USA) and secondary antibody (NA934) was from VWR International (Leicestershire, UK). The intra-assay and inter-assay variations were 4.6% and 10.2%, respectively.

### Validation of AAV injection site

After completion of experiments, mice were anaesthetised with a lethal dose of ketamine and transcardially perfused with heparinised saline for 5 min, followed by 10 min of ice-cold 4% paraformaldehyde (PFA) in phosphate buffer (pH 7.4) for 15 min using a pump (Minipuls, Gilson, Villiers Le Bel, France). Brains were rapidly collected and postfixed sequentially at 4 °C in 15% sucrose in 4% PFA and in 30% sucrose in phosphate-buffered saline until they sank. Afterwards, brains were snap-frozen on dry ice and stored at −80 °C until processing. Brains were coronally sectioned (30-μm) using a cryostat (Bright Instrument Co., Luton, UK) and every third section was collected between −1.34 mm to −2.70 mm from the bregma. Sections were mounted on microscope slides, air-dried and cover slipped with ProLong Antifade mounting medium (Molecular Probes, Inc. OR, USA). The injection site was verified and evaluated and only animals expressing mCherry fluorescent protein in the MePD were included in the analysis by using Axioskop 2 Plus microscope equipped with Axiovision, version 4.7 (Zeiss).

### LH Pulses and Statistical Analysis

Detection of LH pulses was established by use of the Dynpeak algorithm (15). The effect of optogenetic stimulation on parameters of LH secretion was calculated by comparing the mean LH inter-pulse interval within the 90 min stimulation period with the 60 min pre-stimulation control period. For the non-stimulated control animals, the same timepoints were compared. The mean interval between LH pulses, within the 90 min stimulation period, or equivalent, was also compared between experimental groups. Statistical significance was tested using a two-tailed paired t-test. P < 0.05 was considered statistically significant. Data are presented as the mean ± SEM.

### Immunohistochemistry

Mice were anesthetized with an overdose of pentobarbital (Euthatal, *Merial* Animal Health Limited. UK) and perfused transcardially with 15 ml of 4% paraformaldehyde in 0.1 M phosphate buffered saline pH 7.6. (Sigma-Aldrich, UK). The brains were removed and post-fixed in the same fixative at room temperature for 1 hour and then transferred to cryoprotectant media (sucrose 30%, polyethylene glycol 30%, polyvinylpyrolidone 1% in 0.1M phosphate buffer, pH 7.2) until sectioning. The brains were cut on a microtome with a freezing stage into 60 µm coronal sections and stored in cryoprotectant media at −20 °C.

Free-floating sections were washed in Tris-buffered saline (TBS, pH 7.5) three times to remove the cryoprotectant media, then subjected to antigen retrieval in citrate buffer (pH 6.0) at 80 °C for 30 min. The sections were washed in TBS three times then blocked with 5% BSA (Sigma-Aldrich, UK) and 0.5% Triton X (Sigma-Aldrich, UK) for 15 min at room temperature on an orbital shaker. Sections were incubated with rabbit anti-Kisspeptin (AC566,1:2000 dilution; generous gift from Alain Caraty, France) or rabbit anti-GABA (A2052,1:600 dilution, Sigma-Aldrich, UK) in 2% normal goat serum (NGS; Sigma-Aldrich, UK) and incubation media (0.05 M Tris pH7.5, 0.15 M sodium chloride with 0.25% Triton-X-100, 0.3% bovine serum albumin, 2% normal serum, buffered to pH 7.6) for 48 h at 4 °C on an orbital shaker. After the primary antibody incubation, sections were washed three times in TBS and then incubated with biotinylated goat anti-rabbit secondary antibody (1:200 dilution in incubation media; Vector Laboratories, USA) for 2 h at room temperature on an orbital shaker. Sections were washed three times in TBS and then incubated with fluorophore 488-conjugated biotinylated goat anti-rabbit (1:400 dilution; Thermo Fisher Scientific, UK) for 2 h at room temperature in the dark on an orbital shaker. Sections were then washed three times in TBS and mounted on poly-lysine-coated slides (VWR, UK) and stored at 4 °C in the dark until being imaged using epi-fluorescence and confocal microscopy (Leica SP2 Laser Scanning Confocal Microscope, Cambridge Advanced Imaging Centre, UK).

## Results

### Validation of AAV injection site

The AAV-ChR2 virus used to infect the cells in Kiss-CRE mice was tagged with fluorescent mCherry, allowing it to be visualised under a microscope. Analysis of images acquired from coronal sectioning of the mouse brains showed that 6 out of the 8 animals had successful stereotaxic injection of AAV-ChR2 virus into the MePD. The mean number of mCherry-positive cells in unilaterally-injected brain sections was 22.00 ± 4.81. A representative example of a coronal brain section is presented in figure 1. Animals with misplaced injections did not have altered LH pulse frequencies following optogenetic stimulation (data not shown).

**Figure 1:**
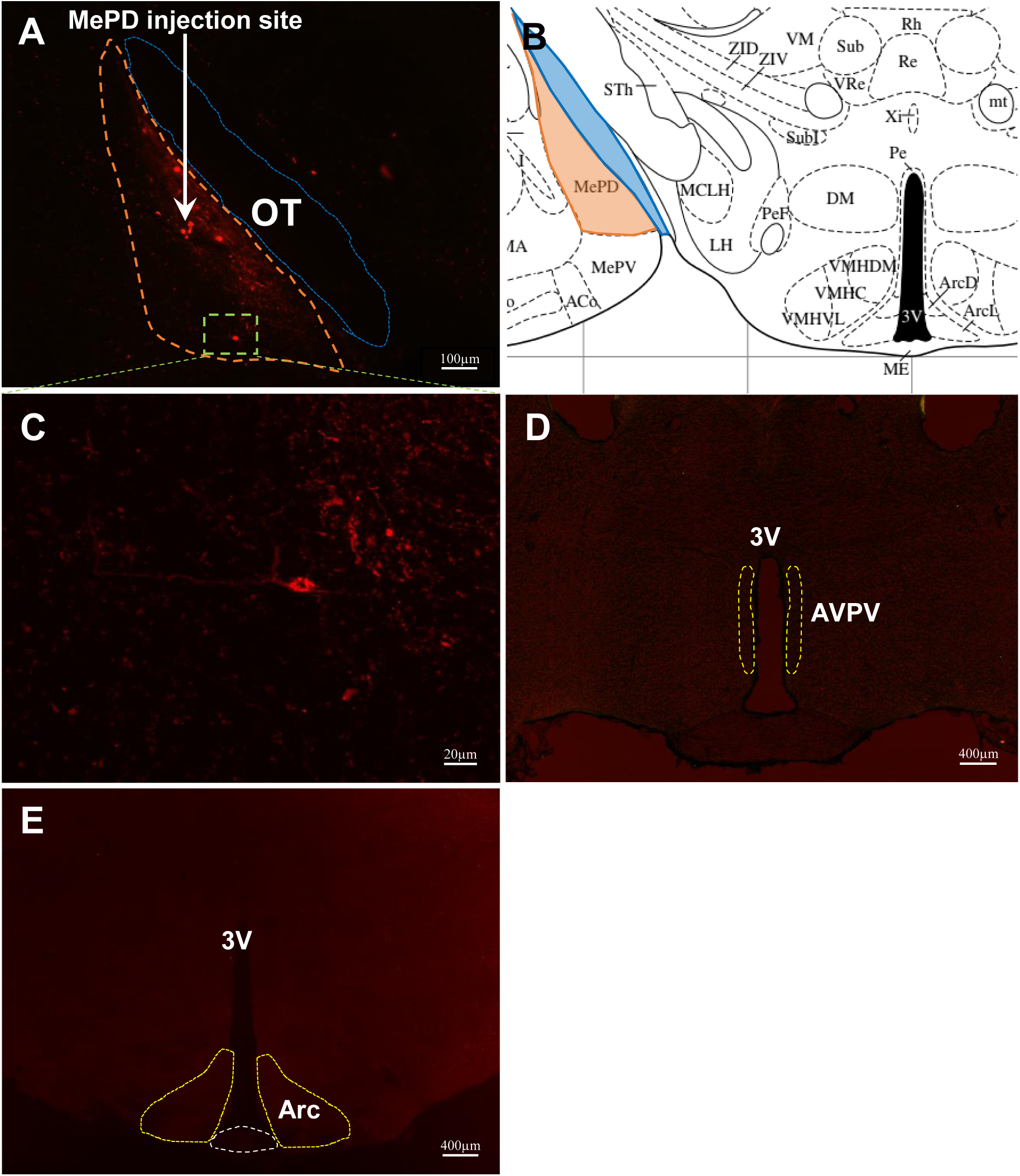
Expression of MePD kisspeptin neurones with ChR2-mCherry in Kiss-CRE mice. Coronal section showing red mCherry fluorescence positive neurones (orange line) in the MePD (A) and the white arrow indicates the injection site of AAV9.EF1.dflox.hChR2(H134R)-mCherry.WPRE.hGH into the MePD of Kiss-Cre mice in the same section. Higher-power view shows the MePD kisspeptin neurones tagged with mCherry (red fluorescence), which indicates ChR2 receptor expressing kisspeptin neurones (C). The absence of mCherry fluorescence in the AVPV (D) and ARC or median eminence (ME) (E) shows the specific infection of MePD kisspeptin cells with the channelrhodopsin. Schematic representation of MePD and its spatial relationship with the optic tract (B). ME, median eminence; OT, optic tract; 3V, third ventricle.

### Effects of continuous optogenetic stimulation of MePD kisspeptin neurones on LH pulse frequency

Following 1 h blood sampling with no light stimulation, MePD kisspeptin neurones were stimulated with blue light at 0.5, 2, or 5 Hz for 90 min. In Kiss1-CRE mice, only continuous stimulation at 5 Hz resulted in a significant increase in LH pulse frequency from a mean inter-pulse interval (IPI) of 26.07 ± 3.81 min to 16.64 ± 0.83 min (Fig. 2). No significant change in IPI was observed at 0.5 or 2 Hz optic stimulation, or in the non-stimulated controls (Fig. 2). Additionally, in wild-type Kiss1-CRE mice, there was no change in LH pulse frequency at 5 Hz optic stimulation (data not shown).

**Figure 2:**
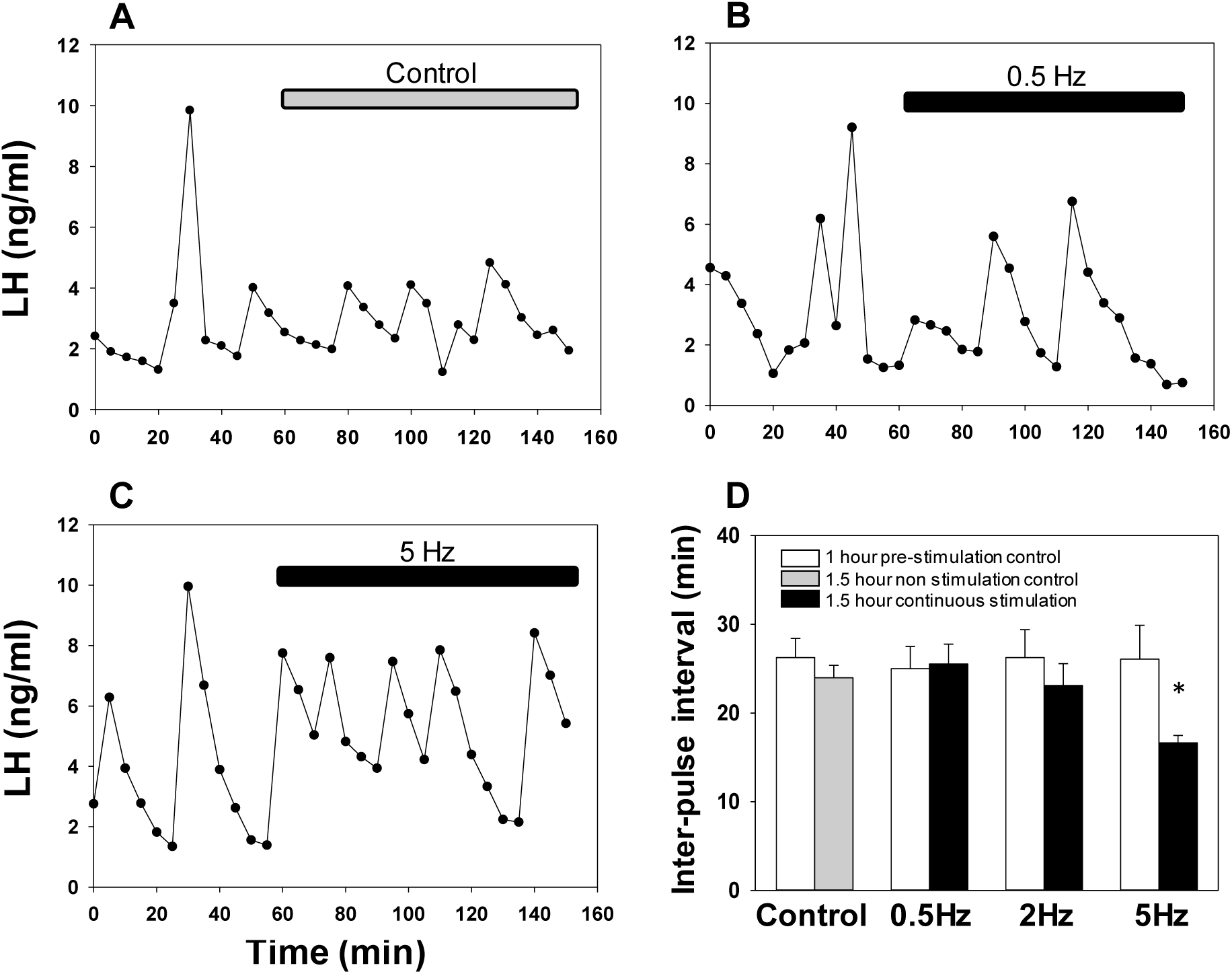
Effect of continuous optogenetic stimulation of MePD *Kiss1* neurones on LH pulse frequency. Representative examples showing the effects of no stimulation (A), 0.5 Hz (B), or 5 Hz stimulation (C) on pulsatile LH secretion in ovariectomised mice. (D), The mean LH inter-pulse interval for increasing frequencies before and after the onset of stimulation. Continuous stimulation at 5 Hz resulted in a significant reduction in the LH inter-pulse interval *P < 0.05 vs control, (n = 6 per treatment group). Results represent mean ± SEM.

### *Kiss1* neurons in the medial amygdala do not co-express GABA

GABA expression was analysed by immunohistochemistry in tdTomato positive *Kiss1* neurons in both the AVPV (Fig. 3A) and the MePD (Fig. 3B). In the AVPV region, we found that 14/20 *Kiss1* neurons, co-expressed GABA but in the MePD, no co-expression was found (n=2 mice, n=8 neurons).

**Figure 3:**
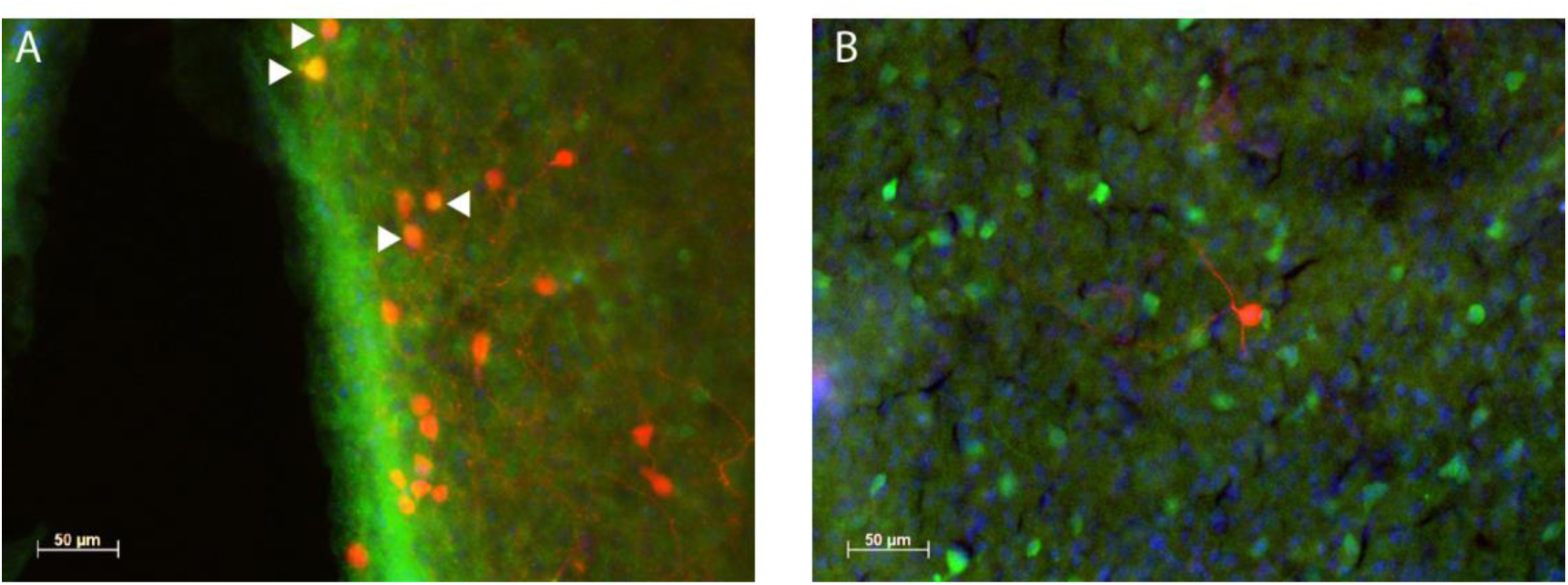
GABA expression in Kiss1-tdTomato labelled neurones. (A). AVPV region. Arrowheads indicate Kiss1-tdTomato neurones (red) that co-express GABA (orange). (B). MePD region showing a single Kiss1-tdTomato neurone (red) that does not co-express GABA (green). Nuclei were counterstained with DAPI (blue).

## Discussion

The present study demonstrates that optogenetic stimulation of *Kiss1* neurones within the MePD increases LH pulse frequency and provides clear evidence that MePD *Kiss1* neuronal activity can modulate the frequency of the hypothalamic GnRH pulse generator. The relationship between the amygdala and its regulation of gonadotrophic hormone secretion has received considerable attention, with evidence to support the notion that the medial amygdala in fact has a negative output on the reproductive system. Stimulating the medial amygdala results in delayed pubertal onset (16), whilst lesions advance the onset of puberty (17) and increase the secretion of LH (18). Indeed, the findings that up to 70% of the neuronal outputs arising from the MePD are GABAergic (19), including a significant percentage of those reaching reproductive centres (20), support the idea of an MePD inhibitory modulation of reproduction. We have unpublished observations showing MePD neurones projecting directly onto KNDy neurones and forming synaptic inputs (21) although their neurochemical phenotype remains unknown. Nevertheless, identification of the latter connection to the KNDy network might help to establish the underlying mechanisms by which the MePD regulates the GnRH pulse generator. Interestingly, MePD *Kiss1* neurones have been shown to project to GnRH neurones in the medial preoptic area (POA) (22). However, our previous data showing Kiss1 receptor antagonism in the MePD reduced LH pulse frequency (10) would suggest that MePD Kiss1 signalling *per se* rather than Kiss1 output to the POA GnRH neurones underlies the upstream regulation of the hypothalamic GnRH pulse generator. Our present finding that continuous lower-frequency optic stimulation increased the frequency of LH pulses supports the concept that MePD control of LH secretion is via activation of the KNDy neurones in the ARC, which are thought to be a major component of the neural construct underlying GnRH pulse generation (3).

How do we reconcile the conundrum that classical stimulation and lesion studies suggest an inhibitory influence of the amygdala, whilst selective optic stimulation of MePD Kiss1 enhances LH secretion? Although GABAergic neurones are present in the MePD, we have not found co-localisation between GABA and tdTomato labelled *Kiss1* neurons in this region. The specificity of the anti-GABA antibody was confirmed by showing around 70% co-localization between GABA and *Kiss1* tdTomato neurones in the AVPV region consistent with data reported elsewhere (23). Thus, specific activation of *Kiss1* neurones in the MePD would not be expected to be inhibitory on LH secretion. An alternative, but less simple explanation might involve a disinhibitory system whereby Kiss1 in the MePD activates inhibitory GABAergic interneurons, which in turn inhibit GABAergic efferents to the ARC. Whilst this remains hypothetical, it is important to realise that the MePD is of pallidal origin (24), therefore suggesting that a neural circuit within this structure is, at least in part, of a classical GABAergic disinhibitory nature. Although the endogenous firing pattern of MePD Kiss1 neurones is unknown, we have shown that continuous optogenetic activation of this population at the relatively low frequency of 5 Hz increased LH pulse frequency, thus demonstrating the minimal activational requirements to modulate the hypothalamic GnRH pulse generator. Further work is required to establish the functional link between MePD *Kiss1* neurone activation and GABAergic signalling to affect pulsatile LH secretion.

Evidence that *Kiss1* neurones within the MePD have a stimulatory role modulating the GnRH pulse generator is an important development in reproductive neuroendocrinology, as it provides further mechanistic insight into why KISS1R receptor antagonism in the MePD was shown to delay puberty, disrupt oestrous cyclicity and reduce the occurrence of the preovulatory LH surge in the rat (8). The MeA is highly-active in response to a range of external stressors such as restraint (25), footshock (26), and predatory odour (27), and is involved in the activation of the hypothalamic-pituitary-adrenal (HPA) axis in response to external anxiogenic stimuli; increased levels of ACTH and glucocorticoids are accompanied with stimulation of the MeA (28). Additionally, as part of the limbic brain, the MePD is an upstream regulator that likely acts as a neural hub integrating several external signals, including olfactory chemosignals and anxiogenic stimuli, with the function of the GnRH pulse generator. Kp and the amygdala have been shown to be critically involved in the integration of olfactory cues with the promotion of sexual behaviour. Lesioning of the MePD results in a diminution of sexual partner preference in rodents (29), and in the “ram effect”, whereby the introduction of a sexually active ram can overcome reproductive inactivity in anoestrous ewes, the resurgence of functional LH pulsatility coincides with an increase in *Kiss1* neurone activity (FOS) in the ARC, POA, and MeA (30, 31). Our previous study using pharmacological activation of MePD *Kiss1* neurones with the DREADD (Designer Receptors Exclusively Activated by Designer Drugs) methodology built upon previous understanding of the facilitatory role of the MePD in sexual behaviour. We showed that stimulating MePD *Kiss1* neurones in this way produces a dual result of augmenting sexual partner preference whilst attenuating anxiety behaviour in male mice (9). To support these findings, we also have unpublished data showing optogenetic stimulation of MePD *Kiss1* neurones leading to a reduction in circulating levels of the stress hormone corticosterone. Furthermore, in studies of men viewing sexual images, intravenous administration of kisspeptin results in increased neuronal activity in the amygdala, detected with fMRI neuroimaging, corresponding to a reduction in negative mood and reduced aversion to sex (32). Taken together, these are important developments in reproductive neuroendocrinology. Investigating kisspeptin in the limbic brain has been able to further our understanding of the elusive neural mechanisms governing reproductive physiology, including pubertal development, as well as the mechanisms by which stress has its inhibitory effects on reproductive capability.

## References

1. Seminara SB, Messager S, Chatzidaki EE, Thresher RR, Acierno JS, Jr., Shagoury JK, Bo-Abbas Y, Kuohung W, Schwinof KM, Hendrick AG, Zahn D, Dixon J, Kaiser UB, Slaugenhaupt SA, Gusella JF, O’Rahilly S, Carlton MB, Crowley WF, Jr., Aparicio SA, Colledge WH. The GPR54 gene as a regulator of puberty. N Engl J Med. 2003; 349(17): 1614–27.

2. de Roux N, Genin E, Carel JC, Matsuda F, Chaussain JL, Milgrom E. Hypogonadotropic hypogonadism due to loss of function of the KiSS1-derived peptide receptor GPR54. Proc Natl Acad Sci U S A. 2003; 100(19): 10972–6.

3. Clarkson J, Han SY, Piet R, McLennan T, Kane GM, Ng J, Porteous RW, Kim JS, Colledge WH, Iremonger KJ, Herbison AE. Definition of the hypothalamic GnRH pulse generator in mice. Proc Natl Acad Sci U S A. 2017; 114(47): E10216–E23.

4. Uenoyama Y, Inoue N, Pheng V, Homma T, Takase K, Yamada S, Ajiki K, Ichikawa M, Okamura H, Maeda KI, Tsukamura H. Ultrastructural evidence of kisspeptin-gonadotrophin-releasing hormone (GnRH) interaction in the median eminence of female rats: implication of axo-axonal regulation of GnRH release. J Neuroendocrinol. 2011; 23(10): 863–70.

5. Qiu J, Nestor CC, Zhang C, Padilla SL, Palmiter RD, Kelly MJ, Ronnekleiv OK. High-frequency stimulation-induced peptide release synchronizes arcuate kisspeptin neurons and excites GnRH neurons. Elife. 2016; 5.

6. Lin Y, Li X, Lupi M, Kinsey-Jones JS, Shao B, Lightman SL, O’Byrne KT. The role of the medial and central amygdala in stress-induced suppression of pulsatile LH secretion in female rats. Endocrinology. 2011; 152(2): 545–55.

7. Kim J, Semaan SJ, Clifton DK, Steiner RA, Dhamija S, Kauffman AS. Regulation of Kiss1 expression by sex steroids in the amygdala of the rat and mouse. Endocrinology. 2011; 152(5): 2020–30.

8. Adekunbi DA, Li XF, Li S, Adegoke OA, Iranloye BO, Morakinyo AO, Lightman SL, Taylor PD, Poston L, O’Byrne KT. Role of amygdala kisspeptin in pubertal timing in female rats. PLoS One. 2017; 12(8): e0183596.

9. Adekunbi DA, Li XF, Lass G, Shetty K, Adegoke OA, Yeo SH, Colledge WH, Lightman SL, O’Byrne KT. Kisspeptin neurones in the posterodorsal medial amygdala modulate sexual partner preference and anxiety in male mice. J Neuroendocrinol. 2018.

10. Comninos AN, Anastasovska J, Sahuri-Arisoylu M, Li X, Li S, Hu M, Jayasena CN, Ghatei MA, Bloom SR, Matthews PM, O’Byrne KT, Bell JD, Dhillo WS. Kisspeptin signaling in the amygdala modulates reproductive hormone secretion. Brain Struct Funct. 2016; 221(4): 2035–47.

11. Yeo SH, Kyle V, Morris PG, Jackman S, Sinnett-Smith LC, Schacker M, Chen C, Colledge WH. Visualisation of Kiss1 Neurone Distribution Using a Kiss1-CRE Transgenic Mouse. J Neuroendocrinol. 2016; 28(11).

12. Paxinos G, Franklin K. The mouse brain in stereotaxic coordinates. Boston: Elsevier Academic Press, 2004.

13. Steyn FJ, Wan Y, Clarkson J, Veldhuis JD, Herbison AE, Chen C. Development of a methodology for and assessment of pulsatile luteinizing hormone secretion in juvenile and adult male mice. Endocrinology. 2013; 154(12): 4939–45.

14. Czieselsky K, Prescott M, Porteous R, Campos P, Clarkson J, Steyn FJ, Campbell RE, Herbison AE. Pulse and Surge Profiles of Luteinizing Hormone Secretion in the Mouse. Endocrinology. 2016; 157(12): 4794–802.

15. Vidal A, Zhang Q, Medigue C, Fabre S, Clement F. DynPeak: an algorithm for pulse detection and frequency analysis in hormonal time series. PLoS One. 2012; 7(7): e39001.

16. Bar-Sela M, Critchlow V. Delayed puberty following electrical stimulation of amygdala in female rats. Am J Physiol. 1966; 211(5): 1103–7.

17. Stephens SB, Raper J, Bachevalier J, Wallen K. Neonatal amygdala lesions advance pubertal timing in female rhesus macaques. Psychoneuroendocrinology. 2015; 51 307–17.

18. Lawton IE, Sawyer CH. Role of amygdala in regulating LH secretion in the adult female rat. Am J Physiol. 1970; 218(3): 622–6.

19. Keshavarzi S, Sullivan RK, Ianno DJ, Sah P. Functional properties and projections of neurons in the medial amygdala. J Neurosci. 2014; 34(26): 8699–715.

20. Choi GB, Dong HW, Murphy AJ, Valenzuela DM, Yancopoulos GD, Swanson LW, Anderson DJ. Lhx6 delineates a pathway mediating innate reproductive behaviors from the amygdala to the hypothalamus. Neuron. 2005; 46(4): 647–60.

21. Yeo SH, Manchisi SH, Colledge WH. Mapping neuronal inputs into Kiss1 neurons in the arcuate. International Congress of Neuroendocrinology, Toronto, Canada, 2018.

22. Pineda R, Plaisier F, Millar RP, Ludwig M. Amygdala Kisspeptin Neurons: Putative Mediators of Olfactory Control of the Gonadotropic Axis. Neuroendocrinology. 2017; 104(3): 223–38.

23. Cravo RM, Margatho LO, Osborne-Lawrence S, Donato J, Jr., Atkin S, Bookout AL, Rovinsky S, Frazao R, Lee CE, Gautron L, Zigman JM, Elias CF. Characterization of Kiss1 neurons using transgenic mouse models. Neuroscience. 2011; 17337–56.

24. Pardo-Bellver C, Cadiz-Moretti B, Novejarque A, Martinez-Garcia F, Lanuza E. Differential efferent projections of the anterior, posteroventral, and posterodorsal subdivisions of the medial amygdala in mice. Front Neuroanat. 2012; 6 33.

25. Ma S, Morilak DA. Norepinephrine release in medial amygdala facilitates activation of the hypothalamic-pituitary-adrenal axis in response to acute immobilisation stress. J Neuroendocrinol. 2005; 17(1): 22–8.

26. Rosen JB, Fanselow MS, Young SL, Sitcoske M, Maren S. Immediate-early gene expression in the amygdala following footshock stress and contextual fear conditioning. Brain Res. 1998; 796(1-2): 132–42.

27. Dielenberg RA, Hunt GE, McGregor IS. “When a rat smells a cat”: the distribution of Fos immunoreactivity in rat brain following exposure to a predatory odor. Neuroscience. 2001; 104(4): 1085–97.

28. Herman JP, Ostrander MM, Mueller NK, Figueiredo H. Limbic system mechanisms of stress regulation: hypothalamo-pituitary-adrenocortical axis. Prog Neuropsychopharmacol Biol Psychiatry. 2005; 29(8): 1201–13.

29. DiBenedictis BT, Ingraham KL, Baum MJ, Cherry JA. Disruption of urinary odor preference and lordosis behavior in female mice given lesions of the medial amygdala. Physiol Behav. 2012; 105(2): 554–9.

30. Fabre-Nys C, Cognie J, Dufourny L, Ghenim M, Martinet S, Lasserre O, Lomet D, Millar RP, Ohkura S, Suetomi Y. The Two Populations of Kisspeptin Neurons Are Involved in the Ram-Induced LH Pulsatile Secretion and LH Surge in Anestrous Ewes. Endocrinology. 2017; 158(11): 3914–28.

31. Gelez H, Fabre-Nys C. Neural pathways involved in the endocrine response of anestrous ewes to the male or its odor. Neuroscience. 2006; 140(3): 791–800.

32. Comninos AN, Wall MB, Demetriou L, Shah AJ, Clarke SA, Narayanaswamy S, Nesbitt A, Izzi-Engbeaya C, Prague JK, Abbara A, Ratnasabapathy R, Salem V, Nijher GM, Jayasena CN, Tanner M, Bassett P, Mehta A, Rabiner EA, Honigsperger C, Silva MR, Brandtzaeg OK, Lundanes E, Wilson SR, Brown RC, Thomas SA, Bloom SR, Dhillo WS. Kisspeptin modulates sexual and emotional brain processing in humans. J Clin Invest. 2017; 127(2): 709–19.

